# DNA bridging explains sub-diffusive movement of chromosomal loci in bacteria

**DOI:** 10.1101/2022.11.18.517049

**Authors:** Srikanth Subramanian, Seán M. Murray

**Affiliations:** Max Planck Institute for Terrestrial Microbiology, Marburg

**Keywords:** Bacterial chromosome, Nucleoid associated proteins, DNA bridging, Sub-diffusive dynamics, Polymer simulations

## Abstract

Chromosomal loci in bacterial cells show a robust sub-diffusive scaling of the mean square displacement, MSD(*τ*) ∼ *τ*^*α*^, with *α* < 0.5. On the other hand, recent experiments have also shown that DNA-bridging Nucleoid Associated Proteins (NAPs) play an important role in chromosome organisation and compaction. Here, using polymer simulations we investigate the role of DNA bridging in determining the dynamics of chromosomal loci. We find that bridging compacts the polymer and reproduces the sub-diffusive elastic dynamics of monomers at timescales shorter than the bridge lifetime. Consistent with this prediction, we measure a higher exponent in a NAP mutant (ΔH-NS) compared to wild-type *E. coli*. Furthermore, bridging can reproduce the rare but ubiquitous rapid movements of chromosomal loci that have been observed in experiments. In our model the scaling exponent defines a relationship between the abundance of bridges and their lifetime. Using this and the observed mobility of chromosomal loci, we predict a lower bound on the average bridge lifetime of around 5 seconds.

**Significance Statement:** The bacterial chromosome exhibits dynamics that cannot be explained by simple polymer models. In particular, the mean square displacement of individual chromosomal loci exhibits a power law scaling with an exponent less than that predicted by polymer theory. Here, we use polymer simulations and experiments to show that DNA bridging by Nucleoid Associated Proteins can explain these anomalous dynamics. Consistent with this, we show that in the absence of the bridging protein H-NS, the scaling exponent increases. Chromosomal loci also display rare rapid movements not explainable by polymer theory, even accounting for the viscoelasticity of the cytoplasm. Our simulations show that bridging can additionally explain this behaviour. Finally, we predict a lower bound on the average bridge lifetime within cells.

---

The diffusive dynamics of chromosomal loci have been characterised *in vivo* in several bacterial species by measuring the scaling exponent *α* of the mean square displacement MSD(*τ*) = ⟨ (***r***(*t* + *τ*) − ***r***(*t*))^2^ ⟩ *τ*^*α*^. However, while polymer theory predicts a sub-diffusive scaling exponent of *α* = 2*ν/*(2*ν* + 1) ≈ 0.54 for a self-avoiding Rouse polymer (*ν* ≈0.588), and *α* = 2*/*3 for a Zimm polymer in a good or theta solvent, irrespective of chain length and topology (1, 2), the values measured for chromosomal loci are consistently less than these estimates across different species, strains and conditions (3–8). Fractional Brownian motion (fBm) of chromosomal loci due to the viscoelastic nature of the cytoplasm has been proposed as a possible explanation for this deviation (3, 9). However, this model cannot reproduce the rare but ubiquitous rapid chromosomal movements (RCMs) made by loci (10) and its predictions are inconsistent with a recent study in which compression of the cell was found to only affect the exponent of chromosomal loci and not cytosolic particles (6). Other mechanisms are therefore required to explain the observed low sub-diffusive exponent.

Nucleoid Associated Proteins (NAPs) are DNA-binding proteins that condense and organise the bacterial nucleoid through bridging, bending and stiffening the DNA (11–13). Recent work using high-throughput chromosome conformation capture (HiC) has investigated how these proteins affect the contact probability between any two chromosomal loci, with different NAPs found to promote either short or long-range contacts (14). The obtained two-point contact probabilities have also been used in polymer models to specify an attractive potential between monomers and thereby make predictions about the organisation of the chromosome within the cell (15–18). However, a bottom-up study of the effect of DNA bridging on bacterial chromosome organisation and dynamics has yet to be performed.

Previous theoretical studies of bridging-like behaviour have been mostly in the context of the unentangled and entangled reversible networks and gels formed by associative polymers (19–21). More recently, at the other extreme of high bridge density, computational models have explored how the resulting globular state can explain the organisation of eukaryotic chromatin (22–29). However, there has been no detailed study of the dynamics induced by bridging in the context of the coiled (non-globular) bacterial chromosome.

Here, in the absence of a dynamical theory, we use polymer simulations to investigate how DNA bridging affects the organisation and dynamics (scaling exponent) of the bacterial chromosome. We confirm that bridging can reduce the scaling exponent of individual monomers below the classic prediction of polymer theory and we characterise the dependence on both the number of bridges and their lifetime, observing a linear relationship between the number of bridges and the compaction of the polymer. Consistent with these results, we show experimentally in *E. coli* that deleting the NAP H-NS results in an increase in the scaling exponent compared to wild type. We also find that bridging produces monomer dynamics that display the same rare, rapid movements (RCMs) as have been observed experimentally, i.e. movements inconsistent with fBm. Finally, we use the experimentally observed mobility of loci to fix an internal timescale in our simulations and thereby predict a lower bound for the average bridge lifetime.

## Results

We model DNA as a self-avoiding linear chain on a periodic cubic lattice [Fig. 1(a)] using the Bond Fluctuation Method (BFM) (30, 31). This model is ergodic, allows a large set of bond angles and reproduces Rouse polymer dynamics. Furthermore, since bond vector lengths can have any of five values, the volume occupied by the polymer depends on its conformation and, as shown below, on the presence of bridges. Taking a polymer of length 400, we choose the lattice dimensions (95 × 95 × 95) to obtain a polymer density that approximately matches that of the *E. coli* chromosome (≈ 1%). We choose a lattice spacing of *h* = 0.0056 μm in order to facilitate a comparison to the observed mean square displacement of a chromosomal locus (see below). Given the density of DNA in an *E. coli* cell, this implies that the polymer represents 800 kb of DNA, which is approximately the size of a chromosomal macrodomain (see Supplementary Information), with each monomer representing 2 kb of DNA.

**Fig. 1.**
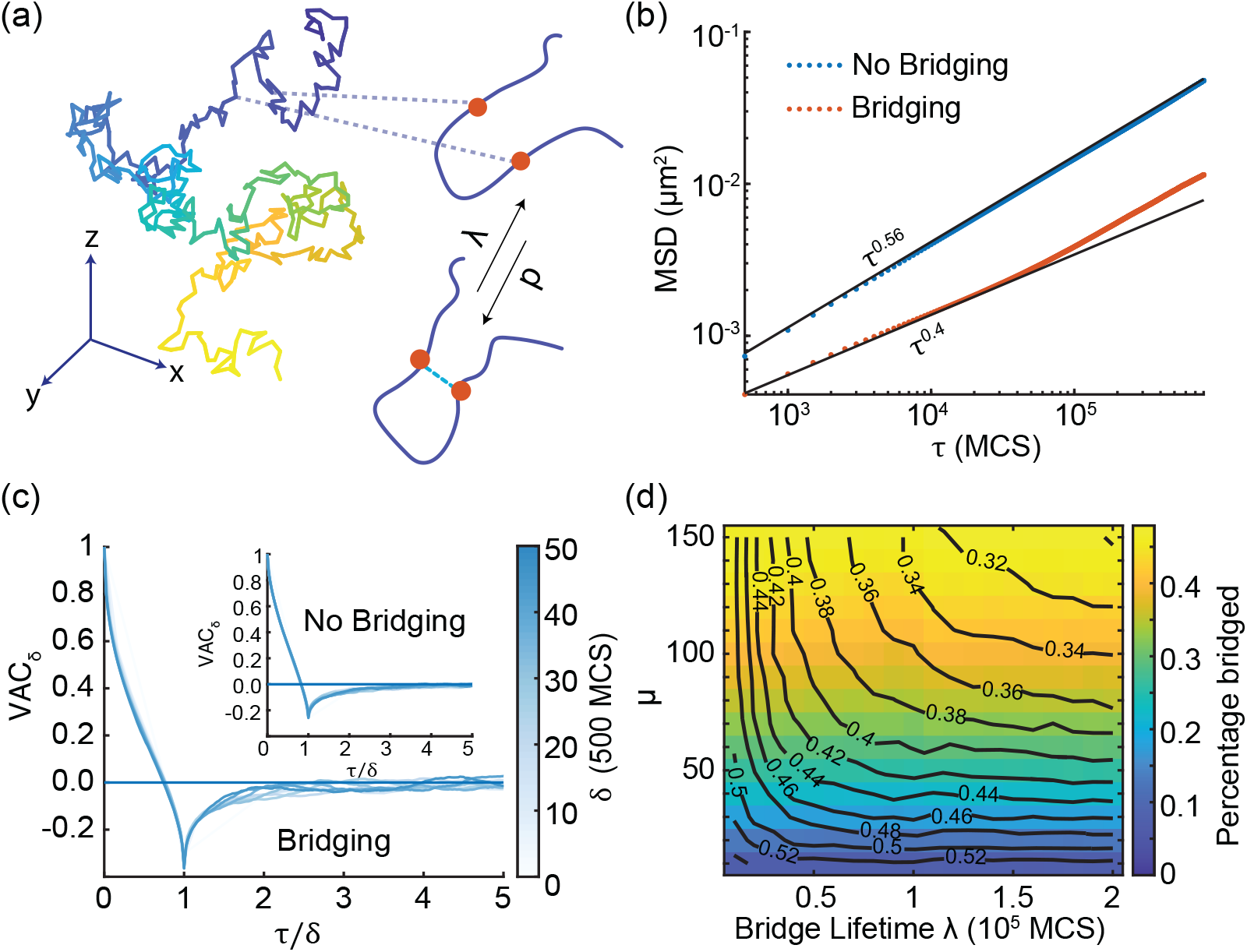
(a) Snapshot of the simulated polymer (N=400). Bridges form between spatially proximal monomers with a probability *p* and have a mean lifetime *λ*. (b) The mean squared displacement (MSD) as a function of delay *τ* with (orange) and without bridging (blue). Bridging reduces the scaling exponent from *α* ≈ 0.56 to *α* ≈ 0.4. Bridging parameters: *p* = 1.5 × 10^*−*3^, *λ* = 4 × 10^4^ MCS. Standard error bars are smaller than the thickness of the lines. (c) Velocity auto-correlation function (VAC) is negative at short lags and collapses for different windows of *τ/δ* indicative of a sub-diffusive process. (d) Phase diagram of the dynamic bridging model in the *μ* =*pλ* and *λ* space. Note that *μ* is positively related to the percentage of monomers bridged. Contours indicate a fixed exponent *α*.

We implement bridging between monomers using the Dynamic Loop model (22, 32). Any two non-neighbouring monomers that are a distance of less than 3 lattice units apart can form a bridge between each other with a probability *p*. Bridges dissociate randomly with probability 1*/λ*, i.e. they have an average lifetime of *λ* in units of Monte Carlo Steps (MCS). While bridged, monomers can still move on the lattice subject to maintaining a bridge length less than 3 lattice units. Each monomer can only form one bridge at a time.

We first confirmed that bridging reduces the scaling exponent of the mean square displacement of a single monomer [Fig. 1(b)], as has been previously shown using the same model in the context of looping of eukaryotic chromatin (22). We found that bridging indeed lowers the exponent: with a level of bridging that results in 28% of monomers bridged, the exponent decreased from *α* ≈0.56 (close to the scaling theory prediction of 0.54) to *α* ≈0.4, a value in line with experimental measurements (3). To examine the dynamics further, we measured the Velocity Auto-Correlation (VAC) function

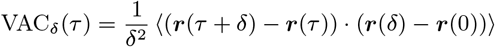

of individual monomers (with the velocity measured over time points *δ* MCS apart). We found that bridging did not change the nature of the VAC [Fig. 1(c)], which remained negative at short time lags with the lowest value at a lag equal to *δ*. This is indicative of elastic or sub-diffusive dynamics and is consistent with experimental measurements of chromosomal loci (33). The VAC has previously been used to distinguish between different models of sub-diffusivity like Fractional Brownian Motion (fBm) and Continuous Time Random Walk (CTRW) (3, 9).

To systematically examine the effect of bridging, we varied the bridging probability *p* and bridge lifetime *λ*, and measured the percentage of monomers bridged and the resultant scaling exponent *α*. We found that the percentage of bridged monomers does not depend on the bridge lifetime individually but only on the product *μ* = *pλ* [Fig. 1(d)]. However, as a dynamical measure, the scaling exponent depends on both *μ* and *λ*; the more bridges and the longer their lifetime, the greater the reduction of the scaling exponent. However, very short-lived bridges (*λ* ≲ S 10^4^ MCS) have little effect on the exponent.

### Bridging compacts the polymer

We next examined the effect of bridging on the organization of the polymer and found a clear effect on compaction [Fig. 2(a)]. This could be quantified using the radius of gyration 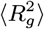 [Fig. S1(a,b)]. However, we found 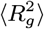 to be a relatively noisy measure of polymer size, motivating us to look for an alternative. The volume occupied by the polymer (*V*) (this is not simply related to the number of monomers since in the bond fluctuation method the excluded volume of different monomers can overlap) was found to be a much more robust measure [Fig. S1(c,d)]. Interestingly it showed a clear linear decrease with the number of bridges formed [Fig. 2(b)]. This was independent of the bridge lifetime *λ* with curves of different *λ* collapsing onto the same line. The latter confirms results from previous Brownian dynamics simulations that the polymer relaxation time can be controlled (though the bridge lifetime) independently of the equilibrium structure (34). The compaction could also be seen by the reduction in the mesh size (from 120 nm without bridging to 85 nm with bridging) [Fig. S1(f)]. Note that, at the levels of bridging studied, the polymer does not enter the globule regime as evidenced by the scaling exponent of the distance between two monomers and the contact probability [Fig. S1(g,h)]. This regime has been studied elsewhere (22, 28, 34).

**Fig. 2.**
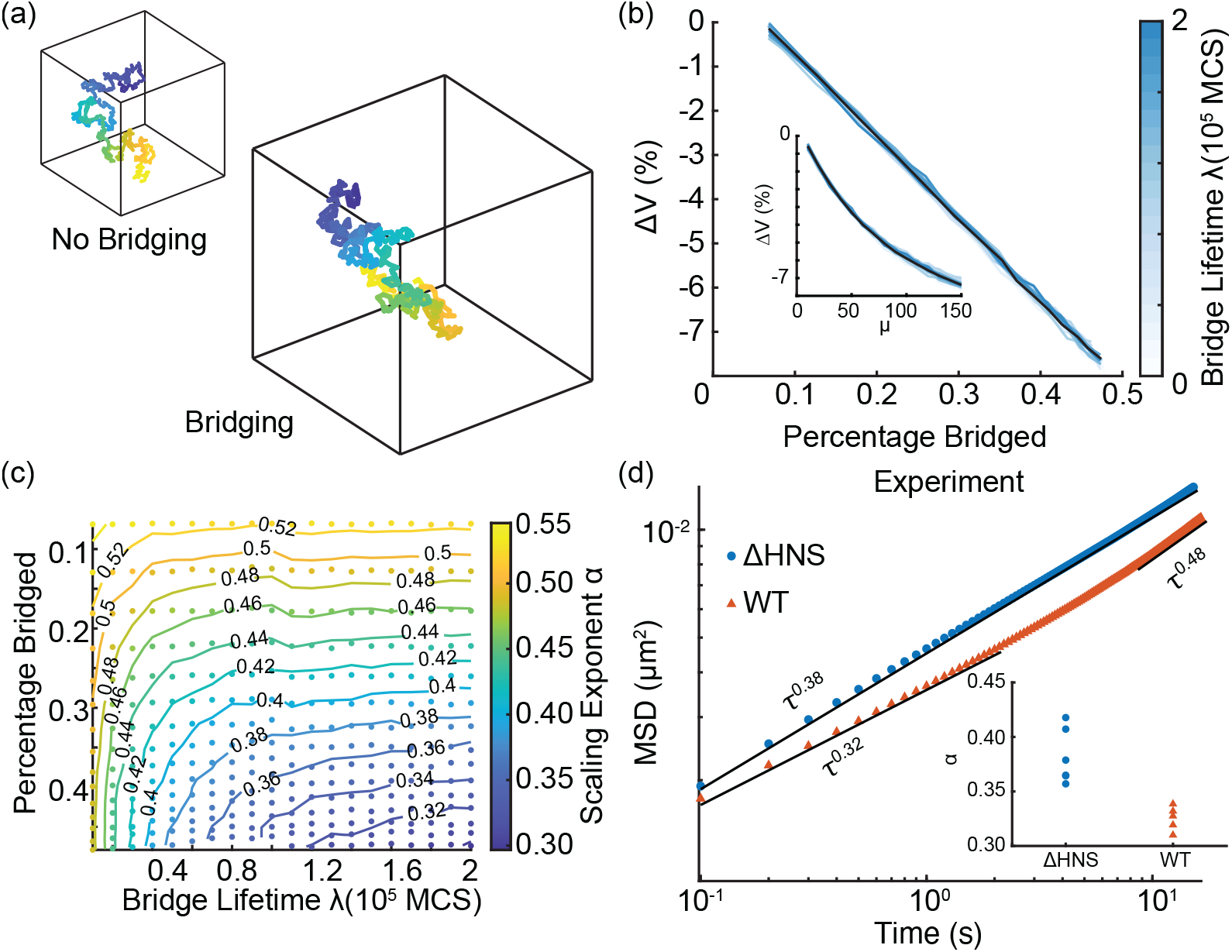
(a) Example polymer conformations with and without bridging. (b) The change in volume occupied relative to the non-bridging polymer Δ*V* decreases linearly with bridging. (Inset) The relationship between Δ*V* and the model parameter *μ*. Black line indicates the average across bridge lifetimes. (c) Phase diagram from Figure 1(d) with *μ* remapped to the percentage of monomers bridged. (d) Loci tracking experiments. Ensemble averaged MSD curves of wild-type *E*.*coli* and a strain deleted of the NAP H-NS. The deletion strain has a higher scaling exponent (*α* ∼ 0.38) compared to the WT (*α* ∼ 0.32). MSD curves are fitted up to a 2 s delay. ΔH-NS: 18089 tracks, WT: 6717 tracks. Inset: Exponent *α* from individual replicates (see Supplementary Information).

Thus far, our simulation parameters were chosen to match the density of DNA within an *E. coli* cell rather than the effect of confinement due to the cell boundaries since the former quantity is likely to more strongly affect the probability of bridge formation. Due to the different scaling of these two effects (see Supplementary Information), a coarse-grained model cannot match both of them simultaneously. Nevertheless, since the *E. coli* chromosome is confined within the cell (filling the cytosol and expanding with cell growth (7))), we also examined how confinement and cuboidal geometry affects bridging [Fig. S2, S3]. The results were qualitatively the same as shown above. However, we note that the mesh size decreased further to the experimentally measured value of 50 nm (35).

To obtain more meaningful physical insights we next remapped the phase diagram in terms of the average number (percentage) of monomers bridged rather than the parameter *μ* [Fig. 2(c)]. This makes it clear that, for sufficiently long bridge lifetimes, an increase (decrease) in the number of bridges formed is concomitant with a decrease (increase) of the scaling exponent. As discussed above, Nucleoid Associated proteins (NAPs) bridge and compact the chromosome. Therefore, our model predicts that the deletion of a bridging NAP should increase the scaling exponent. To test this prediction we measured the MSD scaling exponent of a chromosomal locus in a strain lacking the NAP H-NS. We choose this protein as its deletion has a very mild phenotypic effect compared to other NAPs like HU or MukBEF (14). Following previous work (5), we used fluorescence microscopy and a GFP-ParB/*parS* labelling system to track the *ori* locus of *E. coli* on short timescales. The ensemble-averaged MSD of both the wild type and the ΔH-NS strain are shown in Figure 2(d). Consistent with our model, we found that the scaling exponent *α* for the ΔH-NS (*α* ∼ 0.38) is greater than that of the wild type *α* ∼ 0.32, consistent with a decrease in the number of DNA bridges. While, we observe some variability in the exact value of the exponents between biological replicates (see inset), the exponent of ΔH-NS is consistently higher than the wild type (see also Supplementary Material). This result was not attributable to differences in the signal intensity between the strains (Fig. S10). We conclude that the bridging of chromosomal DNA by nucleoid associated proteins affects the nature of chromosome dynamics and can explain why the scaling exponent of chromosomal loci is more sub-diffusive than expected from polymer dynamics alone.

### Bridging reproduces rapid chromosomal movements

A previous analysis of chromosomal loci dynamics identified a sub-population of fast moving trajectories that could be not be explained by the null phenomenological model of fractional Brownian motion (10). Instead, these outliers, termed Rapid chromosomal movements (RCMs), were speculated to be due to an active machinery or some stress-relaxation mechanism (10, 36). To determine if RCMs were also present in our data, we followed the same procedure as Javer *et al*. to identify them.

We fit the wild type ensemble averaged MSD curves to a power law 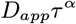to determine the two parameters of the fBm model (37), namely the apparent diffusion constant *D*_*app*_ and the exponent *α* (Fig. 3(a)). These parameters were then used to simulate fBm using the same number and track lengths as the experimental data. We then calculated the drift velocity *v*_*d*_ for each track, defined as the magnitude of the displacement along the major axis between two time points divided by the elapsed time. Similar to Javer *et al*., we found that the distribution of drift velocities of the experimental data displayed a fatter tail than that of the fBm simulations (Figure 3(b)). This disparity was not dependent on the precise elapsed time used nor on the deviation of the MSD curve from a perfect power law (or the range over which the parameter fitting was performed). Indeed, we found that RCMs were also present in the ΔH-NS strain, which displays a near perfect power law behaviour (Fig. S6 (a,b)).

**Fig. 3.**
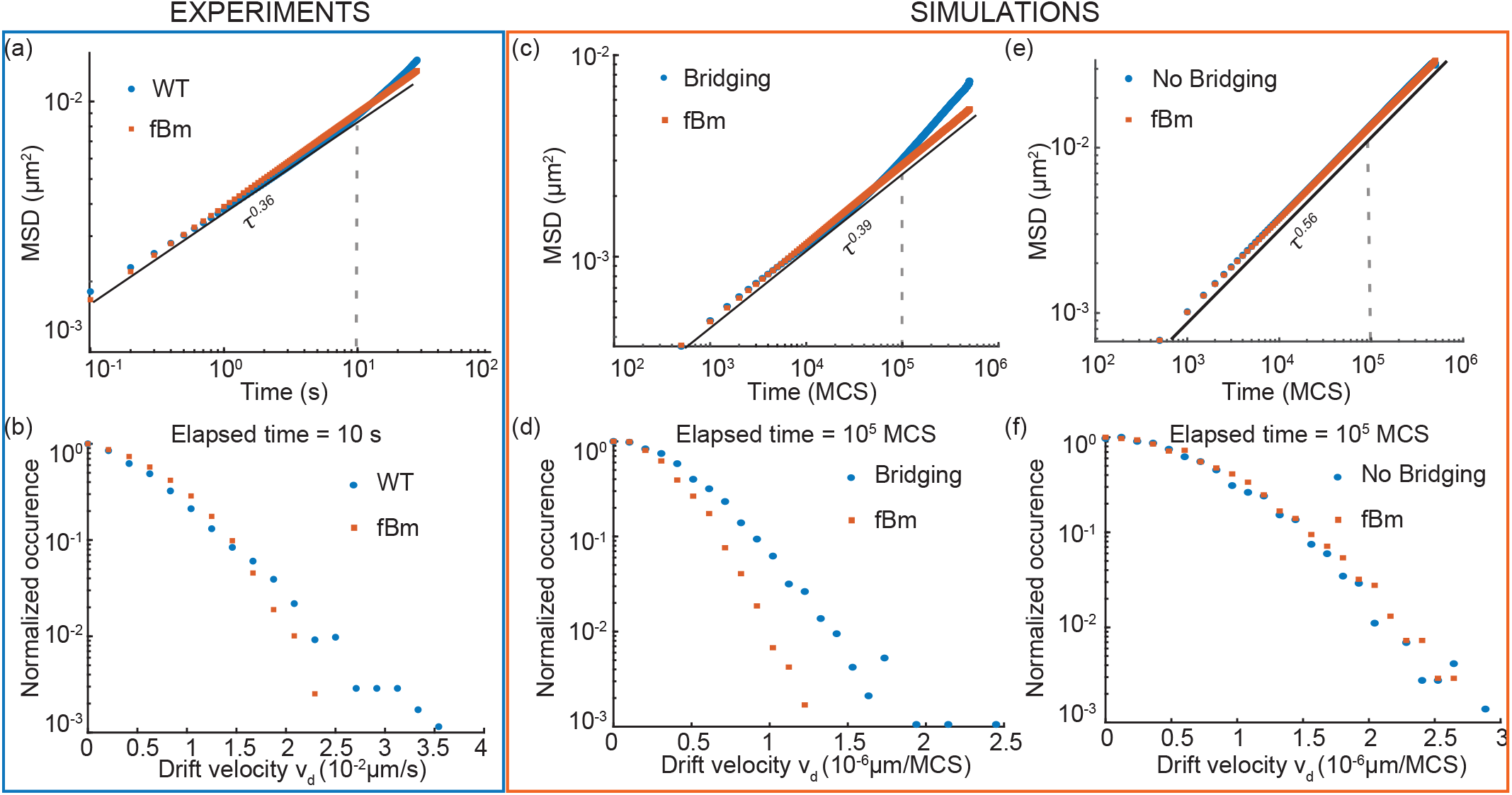
(a) Ensemble averaged MSD from experimental data of WT (blue circles) is fitted to fBm (orange squares, Fit 0.0008*τ* ^0.36^*μm*^2^, 6717 tracks). (b) Drift velocity of wild type tracks (blue circles) have a much wider tail than from simulations of fBm with model parameters chosen according to the fit in (a) (orange squares). (c) Ensemble averaged MSD from polymer simulations fitted with fBm simulations (orange squares, *μ* = 80, *τ* = 10^5^ MCS, Fit 0.0009*τ* ^0.39^*μm*^2^, 5200 tracks). (d) Drift velocity distributions *v*_*d*_ of fBm (orange square) have a smaller distribution than bridging tracks (blue circles). (e) Same as (a,c) but to polymers with no bridging. (f) The distributions of *v*_*d*_ overlap.

Following the same procedure, we then parameter-matched the fBm model using the ensemble averaged MSD curves of our simulated monomer trajectories (Fig. 3(c)). Surprisingly, we found that the bridging simulations produced trajectories with a similar over-representation of high drift velocities compared to the fBm model as seen for the experimental data (Fig. 3(d)). We could directly attribute this disparity to the effect of bridging since simulations without bridging showed no such disagreement (Fig. 3(e,f)). Again, these results persisted irrespective of the precise fitting to the MSD curves and the elapsed time used (Fig. S4,S5,S6). Note that, while bridging produces these outlier movements, overall it slows the dynamics of the polymer and therefore results in lower drift velocities. Consistent with this, the MSD and drift velocities of ΔH-NS were slightly greater than WT (Fig. 2(d), Fig. S6(c)).

We explain the presence of RCMs in our simulations as being due to the heterogeneity in the bridging state of the tracked monomers. On timescales much longer than the mean bridge lifetime, each segment of the polymer is likely to be bridged for the same percentage of time and the dynamics are therefore relatively homogeneous. However on timescales less than the bridge lifetime, there is greater heterogeneity - some segments will remain bridged throughout, others will remain unbridged. This results in a corresponding variation in the observed dynamics, with more strongly bridged segments displaying more restricted movements and shifting a proportion (the majority at high bridging) of the drift velocity distribution to lower values, leaving the unbridged segments as outliers. This heterogeneity cannot be captured by the fBm model, which models a single population. This interpretation is consistent with our finding that the fraction of RCMs increases with the degree of bridging and as the drift velocity is measured over shorter elapsed times (Fig. S5).

While we have not been able to quantitatively fit our polymer simulations to the experimental MSD curve and drift velocity distribution due to the increasingly computationally challenging of simulating longer bridge lifetimes and the higher number of simulations required to obtain accurate statistics of the RCMs, we nevertheless conclude that bridging by NAPs provides a potential explanation for both the sub-diffusive scaling of chromosomal loci as well the observed rare rapid chromosomal movements.

### Transition to a higher exponent places bounds on bridge lifetime

In our simulations of bridging, we observed that the MSD curve transitions at longer time lags to the exponent expected in the absence of bridging (≈0.56 in our model, lower if we were to account for viscoelasticity) [Fig. 4(a)]. Interestingly, we found a similar upward transition in the experimental MSD curve of *ori* [Fig. 2(d)]. This was also observed in previous studies performed at the same (0.1 s) and longer time resolutions (1 s) and was associated to the RCMs discussed above (5, 6, 10). While the cause of this transition is unknown and confounding effects cannot be completely discounted, it is most apparent for the terminus (*ter*) region (5, 6, 10). Since the Ter macrodomain is affected by NAPs (MukBEF in particular) differently than the rest of chromosome (resulting in a bias towards short-range genomic contacts) (14), this is consistent with bridging by NAPs being ultimately responsible. In our simulations, the transition is due to bridges not having an effect on the dynamics at timescales much longer than the bridge lifetime. Indeed, we observe a linear relationship between the transition location and the bridge lifetime *λ* [Fig. 4(b)]. We also note that the transition was not visible in the MSD curve of ΔH-NS strain (at least within the measured range) [Fig. 2(d)], which could be explained by this strain having a longer average bridge lifetime.

**Fig. 4.**
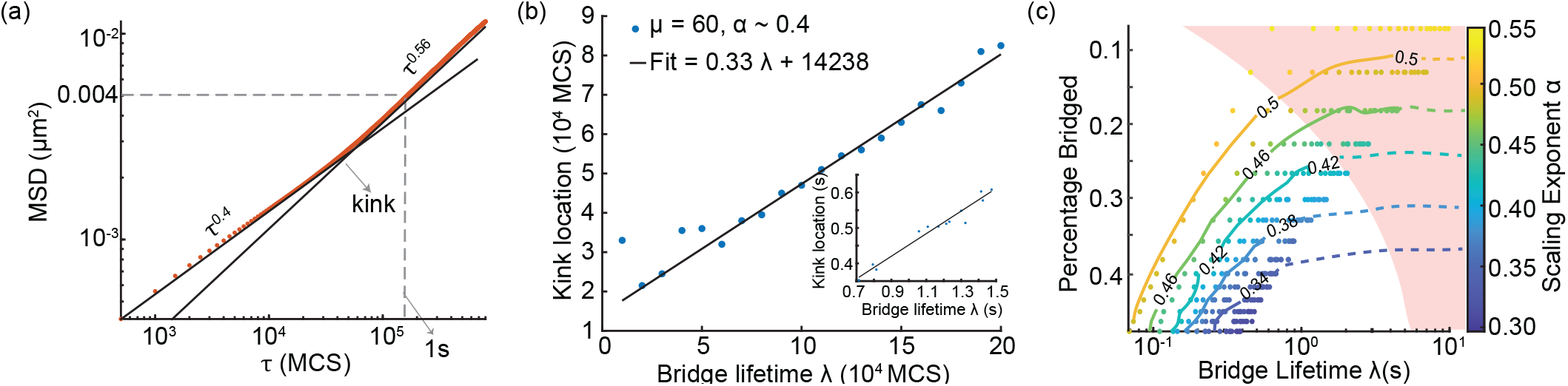
(a) Transition to a higher exponent. MSD curve transitions (at the ‘kink’ point) to a higher exponent *α* ∼ 0.56. (b) Location of the kink depends linearly on the bridge lifetime (*λ*). Inset shows the same plot after conversion to seconds. (c) Phase diagram from Figure 2(c) converted to seconds. The contours of *α* are overlaid. Dashed lines are estimated projections to higher values of *λ*. Shading indicates that the kink lies beyond 1 s.

We next wondered whether the location of the transition could be used to infer a bound on the effective bridge life-time. In this direction, we set the internal timescale of our simulations by matching the MSD of *ori* at a lag of 1 s. In particular, we found an MSD at 1 s of ≈ 0.004 μm^2^ [Fig. 2(d)], consistent with previous measurements (5). With a lattice spacing of 0.0056 μm, this MSD was reached at a lag of ≈ 10^5^ MCS [Fig. 4(a)]. We choose this lattice spacing so as to have sufficient spatial resolution at this displacement. By assigning 10^5^ MCS ≈ 1 s, we can then convert our simulation results from MCS to seconds.

Doing this for all points in the phase diagram we obtain a relationship between two physical measures, the equilibrium percentage of monomers bridged and the bridge lifetime *λ* in seconds for different values of the scaling exponent [Fig. 4(c)]. We have extended the contours of fixed exponent to longer lifetimes by hand as it becomes increasingly computationally challenging to access longer bridge lifetimes (in seconds), especially for the lowest exponents, due to the slow dynamics of the polymer (each second corresponds to an increasingly large number of MCS) [Fig. S7(a,b)]. While the scaling exponent can be measured experimentally, the degree of bridging and the effective bridge lifetime are more challenging to quantify. Nevertheless, the relationship between these variables that we have uncovered here, should be useful in interpreting future experimental results and contributes to our understanding of chromosome dynamics.

Returning to the transition in the MSD curve, we can use the linear relationship to the bridge lifetime [Fig. 4(b)] and the conversion from MCS to seconds described above to obtain the location of the transition in seconds at each point in the phase diagram. In particular, we can identify the region of the phase diagram in which the transition occurs beyond a delay of 1 s, as seen in our data and other measurements. This results in the shaded region in Figure 4(c) and provides a lower bound for the bridge lifetime. For *α*∼ 0.32, as observed for the WT in our experiments, we find a lower bound on the effective bridge lifetime of around 5 s, a reasonable estimate given the relative slow dynamics of chromosomal loci. While measurements of bridge lifetimes of the various NAPs are lacking, estimates for H-NS and HU can be taken from the timescale of their recovery after photobleaching (FRAP) which gave 50 s (38) and 1 s (39) respectively.

## Discussion

Our results provide insight into the role of DNA bridging by nucleoid-associated proteins (NAPs) in determining chromosome dynamics and compaction within bacterial cells. We have shown that bridging, at physically plausible levels and lifetimes, can explain the sub-diffusive scaling exponent of bacterial chromosomal loci. In particular, our model predicts that a decrease in bridging leads to an increase in the sub-diffusive exponent *α* and we confirmed this by tracking the *ori* locus of *E. coli* in a strain deleted of the NAP H-NS. We also addressed the upturn in the ensemble averaged MSD curve seen at long lags. Our model displays a similar transition at a timescale of the order of the average bridge lifetime and we obtained a lower bound on the effective bridge lifetime of the *ori* locus of about 5 s.

Bridging can also qualitatively reproduce the rare but ubiquitous rapid chromosomal movements (RCMs) that are observed within experimental trajectories, in contrast to the null phenomenological model of fractional Brownian motion (fBm). The RCMs in our model are due to the rare event of a DNA segment being unbridged by NAPs for long enough that it exhibits an unusually large movement compared to the rest of the slower-moving bridged polymer. This is consistent with the proposal by Javer *et al*. that RCMs arise due to the relaxation of stress caused by the action of bridging proteins and condensins (10, 40). Therefore, our model provides an explanation for both the sub-diffusive scaling exponent of chromosomal loci and the observation of RCMs. Furthermore, in contrast to the hypothesis of a viscoelastic cytoplasm (3), bridging is consistent with recent work showing that cell compression lowers the exponent of chromosomal loci but not that of diffusive particles (6). A lower cell volume increases the density of DNA and therefore increases the rate of bridge formation, lowering the exponent (Fig. S2). More broadly, by characterising the relation between an equilibrium quantity, the number of bridges, and a dynamic quantity, the mean bridge lifetime (*λ*), our framework provides an intuitive parameter landscape for bacterial chromosome dynamics that will help guide future studies.

## Materials and Methods

### Polymer Simulations

We simulate a self-avoiding linear polymer using the Bond Fluctuation method (BFM) (30), in which each monomer is represented by a cube exclusively occupying 8 lattice vertices and there are 108 allowed bond vectors allowing for a large range of angles. We use the open source software LeMonADE (41) modified to add bridging functionality. The basic (without bridging) simulation is specified by the number of monomers *N* and the lattice dimension *L*. We choose *N* = 400 since it becomes increasingly challenging to simulate longer polymers.

#### Volume Occupied

As each monomer is represented by a cube exclusively occupying 8 vertices, the set of lattice sites occupied by the polymer is the union of 3 × 3 × 3 cubes around the monomers. We define this as the *V*_occupied_ by the polymer in the box. Based on this measure, we find that our polymer of length 400 approximately occupies a total of 8678 lattice sites. Hence, each monomer occupies 21.69 lattice sites on average.

We fix the lattice dimension *L* by using this volume measure to match the volume density of chromosome in the cell (∼ 1%)

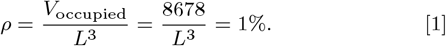

This fixes the dimension of the lattice *L* = 95 lattice units.

In order to compare the simulated MSD data with experiments at short time lags we require sufficient spatial resolution at ∼ 0.004*μm*^2^, the MSD of chromosomal loci at 1 s lag(3, 5). We therefore fix the lattice spacing to be *h* = 0.0056*μm*. The number of base pairs corresponding to a monomer in our simulations is given by,

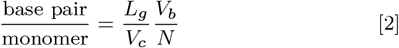

where *V*_c_ ≈ 0.88*μm*^3^ is the volume of the cell, *L*_g_ = 4.5Mbp the length of *E. coli* genome, *V*_*b*_ = *L*^3^*h*^3^ the volume of the box. Hence, we simulate a 800 kb segment (approximately the size of a macrodomain) of the chromosome with each monomer representing a 2kb segment of chromosome. We use periodic boundary conditions in our simulations.

### Monte Carlo procedure

Each simulation is started from a random conformation of the polymer. Monomer diffusion and bridging is implemented in the following manner,

1. Select a monomer at random and attempt a diffusive move (BFM algorithm).
2. Select a random monomer and if co-localised with another free monomer (distance < 3 lattice units), attempt a bridge with probability *p*.
3. Select a random monomer and if bridged, remove the bridge with probability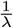.
4. Repeat.

A set of N moves is defined as a single MCS and we sample the configuration every 500 MCS. Note that a bridged monomer can still diffuse as long as the bridge partner is less than 3 lattice units away. We start the Monte Carlo sampling after 5*λ* MCS to ensure sufficient equilibriation of the polymer.

### Calculation of ensemble averaged MSD

The phase diagram is calculated as an average from multiple simulations for each parameter values *μ* = *pλ, λ*. From our simulations, we calculate the ensemble averaged MSD at lag *τ* defined by,

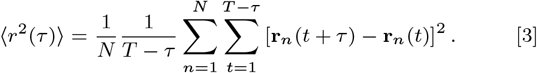

We perform a linear fit to the logarithm of the MSD curves,

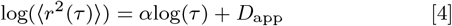

up to a delay of 20000 MCS and obtain ensemble averaged *α*. The data in Fig. 1(b,d) are obtained from 50 (*λ* ≤ 10^5^), 30 (*λ >* 10^5^) independent simulations for each parameter. The ensemble averaged MSD curves are calculated from tracks of every 20th monomer on the polymer (600-1000 tracks per parameter).

### Strains

The *parS/*P1 site from *E*.*coli* strain RM29 obtained from (42) (originally from (43)) was transduced near the *ori* region into MG1655 WT and ΔHNS strains, the latter obtained from the Keio collection of the Sourjik lab (MPI Terrestrial Microbiology). GFP-ParB was expressed from the plasmid pALA2705 with no IPTG induction (43–45). The strains were grown overnight at 30^*°*^C in LB medium with appropriate antibiotics (100*μ*g/mL ampicillin). The overnight culture was diluted into media made of M9–Glucose– Casamino acids (as in (5)) and grown to an optical density of 0.1-0.2.

We chose the P1 labelling system in order to compare our results with previous studies (3, 5, 10, 46). We note that while some differences in the dynamics of the *ter* locus between the ParB labelling systems we use (P1) compared to that of pMT1 have been observed, no substantial differences have been reported for the *ori* locus (8, 44, 45, 47). This agrees with experiments in our lab studying origin positioning and segregation across many thousands of cell cycles.

### Microscopy

1 *μ*L of the sample was placed on 1.5% agraose pads (made of same media as the day culture) and imaged under a Nikon Ti microscope with a 60x/1.4 NA oil objective. The strains were imaged at a constant 30^*°*^C. Images were captured on a Hamamatsu CCD camera using NIS-Elements software. Movies were 450 frames long, with 0.1s interval and an exposure time of 100ms (Fig. S9(a)).

### Analysis

We follow the procedure and analysis in (5). Briefly, foci positions were located via two-dimensional fitting of a Gaussian function to the intensity distributions of individual loci. The ensemble averaged MSD was calculated from pooled trajectories using Eq. (3). The scaling exponent *α* was calculated for ensemble averaged MSD curves by fitting a power law up to 2 s delays.

The codes from Javer et.al was used to track the foci, and are available at https://github.com/ver228/bacteria-loci-tracker. Custom MATLAB scripts were written to analyse the data.

## ACKNOWLEDGMENTS

We thank Ismath Sadhir for help with the strain preparation and Gabriele Malengo for helping set-up the microscopy experiments.

