## Supplementary File for "DNA bridging explains sub-diffusive movement of chromosomal loci in bacteria"

##### This PDF file includes:

Supporting text
Figs. S1 to S10
Table S1
SI References

### Supporting Information Text

#### 1. Equilibrium Properties

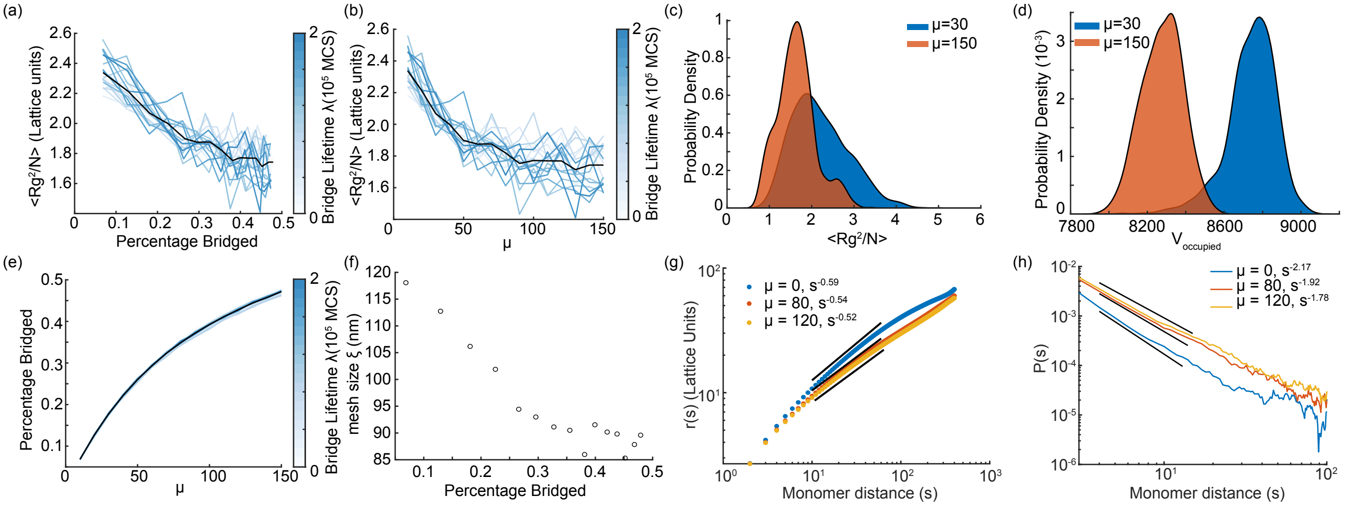


**Fig. S1.** (a) Radius of Gyration $\langle R_g^2/N \rangle$ as function of Percentage Bridged. (b) Radius of Gyration as function plotted as function of $\mu$. (c) Distributions of $\langle R_g^2/N \rangle$ are overlapping for different levels of bridging. (d) Distributions of $V_{\text{occupied}}$ are significantly separated. (e) Percentage of monomers bridged increases with $\mu$ and is independent of bridge lifetime. (f) Mesh size ($\xi$) decreases with increasing bridges. (g) End to end distance for sub-segments on the chain scale as $\langle R \rangle \sim s^{\nu}$. We observe an exponent $\nu = 0.59$ in the absence of bridging as expected for a self-avoiding polymer. The scaling exponent decreases with increasing bridging. But, we do not observe a plateauing of the curves which is indicative of the globule state. (h) The contact probability $P(s)$ between monomers is plotted as function of monomer distance (s). In the absence of bridging (blue curve), we find $P(s) \sim s^{2.18}$ as expected for a self-avoiding chain in a good solvent (1, 2). With increasing bridging the exponent increases, but for our level of bridging we never we are below the compact globule regime.

**$V_{\text{occupied}}$ is robust.** To measure compaction of the polymer we calculated the Radius of Gyration $\langle R_g^2 \rangle$ of the polymer given by,

$$\langle R_g^2 \rangle \equiv \frac{1}{N} \sum_{i=1}^{N} (\mathbf{r}_i - \mathbf{r}_{cm})^2 \qquad [1]$$

as a function of bridging. Radius of gyration decreases with increased bridging but seems to be a noisy measure of compaction (Fig. S1(a),(b)) unlike $V_{\text{occupied}}$ defined in the methods section. In Fig. 2(b) we plot $\Delta V = \frac{V_{\text{occupied}} - V_{\text{no bridging}}}{V_{\text{no bridging}}}$ as a function of bridges and observe linear decrease and a collapse across bridge lifetimes. The distribution of values for parameters across simulations are more distinguishable for $V_{\text{occupied}}$ as compared to $\langle R_g^2 \rangle$ (Fig. S1(c), S1(d)). The ensemble averaged quantities are calculated from $\approx 1000$ independent configurations.

**Mesh size decreases in the presence of bridging.** Here, we calculate the mesh size $\xi$ from our simulations. The probability of finding a monomer at a distance $r$ and $r + dr$ from another randomly chosen monomer is given by $4\pi\rho r^2 g(r)\, dr$, where $\rho = 0.01$ is the density of the polymer and $g(r)$ is a radial density function. For a semi dilute polymer with $r \gg \xi_c$, $g(r)$ is expected to have the form,

$$g(r) = 1 + \frac{A}{r} exp(-r/\xi_c) \qquad [2]$$

where $A > 0$ and $\xi_c > 0$. $\xi_c$ is the correlation length of the polymer which is approximately same as the mesh size $\xi$ for a semi-dilute polymer (3). We calculate $g(r)$ from our simulations and fit to Eq. (2) and find the mesh size ($\xi$) for different parameters. The mesh size of the *E.coli* chromosome was estimated to be around 50nm (4). From our simulations, we find that mesh size decreases with increased bridging and has a range of values between 120-85 nm (Fig. S1(f)).

The End-to-End distance for sub-segments on the chain scale as $r(s) \sim s^{\nu}$ with $\nu \approx 0.588$ (5) for a free polymer (Fig. S1(g)). We observe that with increasing bridging an exponent with decreases, but stays below the globule transition exponent of $\nu = 0.5$. Also, as evidenced by the contact probability curves (Fig. S1(h)) our polymer resides below the globule regime.

**Matching chromosome density versus confinement.** In our simulations, we chose to match the density of *E. coli* chromosome in the cell and not its confinement (average diameter of the polymer relative to the dimension of the box $\frac{2R_g}{L}$). The density $d$ scales linearly with number of monomers $N$, while confinement scales as $R_g \sim N^{\nu}$, where $\nu = 0.588$ making it impossible to match both quantities at the same time. If we fix the confinement of the chromosome instead of its density, the general results in the main text remain unaffected, albeit changing the exact values of $\alpha$ for different parameters. As confinement increases, density increases and $\alpha$ decreases even in the absence of bridging up to $\alpha \approx 0.5$ at which point confinement screens the effect of self-avoidance (Fig. S1)(6).

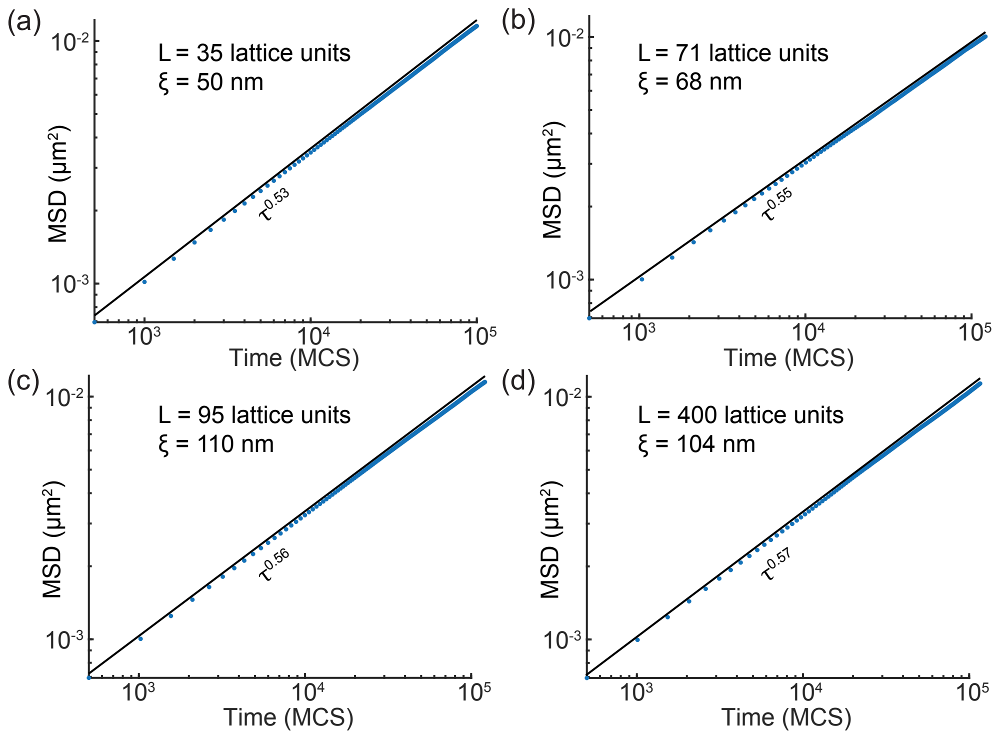


**Fig. S2.** (a)-(d) Confinement affects the scaling exponent of MSD even in the absence of bridging. Mesh size decreases with increasing confinement. Parameters.Lattice size L, Mesh size $\xi$, Polymer length N=400.

Density of the chromosome is clearly the limiting factor for bridging, as a change in density affects the probability of co-localisation for any two monomers. As matching confinement will lead to higher density of the polymer in the box, we matched the density of the chromosome to model realistic effects of bridging by NAPs. Recently, it was observed that the unconstrained *E. coli* chromosome is approximately 2.35 times longer than the cell length (7). By decreasing the lattice size to match the confinement of the *E.coli* chromosome instead of its density we further decrease the mesh size and can match the experimental value of 50nm (see Fig. S1(a)).

Note that increasing confinement increases the number of bridges and consequently decreases the scaling exponent for the same set of parameters $\mu, \lambda$. Importantly, this is consistent with recent work showing that cell compression lowers the MSD scaling exponent of chromosomal loci but not that of diffusive particles (8). A lower cell volume increases the confinement and hence increases the probability of bridge formation of DNA therefore, lowering the exponent.

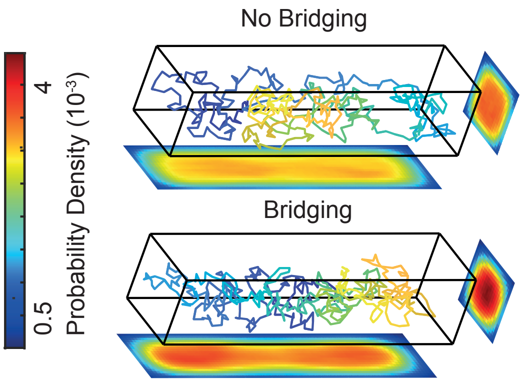


**Fig. S3.** A circular polymer in a cuboid with hard walls matching the confinement of *E.coli* cell. In the presence of bridging the polymer is compacted and has a density that decreases across the cross section. Lattice dimensions $88 \times 22 \times 22$.

*Circular Polymer.—* To illustrate the effect of increased confinement we simulate a circular polymer length $N = 440$ monomers in a box with dimensions $L_x = 88, L_y = 22, L_z = 22$ matching the 4:1 aspect ratio of *E.coli* cell (Fig. S3)). The probability density is calculated by averaging across 1000 independent configurations. In the case of the polymer with no bridges, we observe a uniform distribution of monomer density. In contrast, the bridging polymer shows a radially decreasing density across the cross section. This is notable, as direct imaging of an abundant NAP HU in *E.coli* also revealed a similar decreasing radial density of the chromosome across the cross-section of the cell (9, 10).

#### 2. Bridging reproduces RCMs

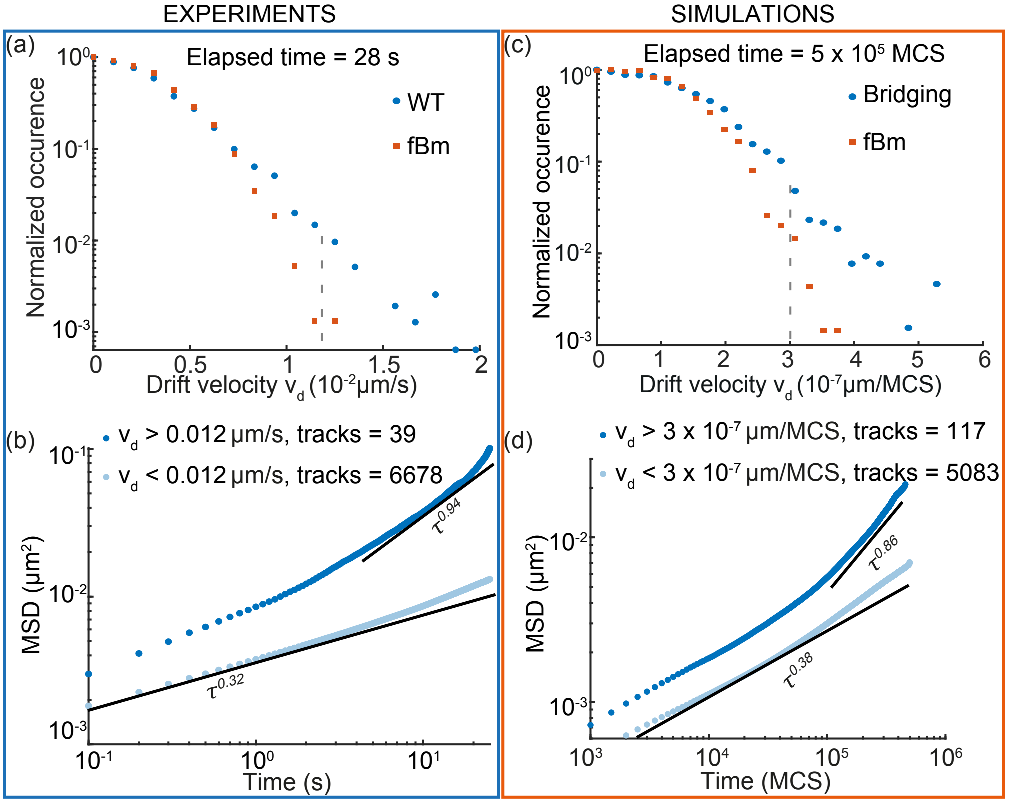


**Fig. S4.** (a) Drift velocity distribution comparisons between WT data and fBm measured over 28s (entire track length). (b) Ensemble averaged MSD of the wild type strain with tracks selected based on drift velocity $v_d$ (grey dashed line in (a) shows the $v_d$ threshold). Tracks with $v_d$ greater that fBm distribution have a transition to higher exponent at longer time lags. (c) Same as in (a) for the bridging simulations with $v_d$ defined over the entire track length of $5 \times 10^5$ MCS. (d) We find a similar subset of tracks (split based $v_d$, grey dashed line in (c)) which transition to higher exponent at longer time lags. Parameters $\mu = 80, \lambda = 10^5$ MCS.

In our experimental data, we select a subset of outlier trajectories (39 of 6717 tracks) with drift velocity $v_d > 0.012\mu ms^{-1}$ (Fig. S4(a)). The ensemble averaged MSD curves show a transition to faster dynamics at longer timescales (see Fig. S4(b)) indicating the presence of RCMs (11). Strikingly, selecting a similar subset of faster tracks (117 of 5083 tracks) with $v_d > 3 \times 10^{-7}\mu m\text{MCS}^{-1}$ not captured by fBm (Fig. S4(c)) in our bridging simulations, we find similar MSD curves. The tracks with faster $v_d$ transition to a higher exponent at longer time lags (Fig. S4(d)).

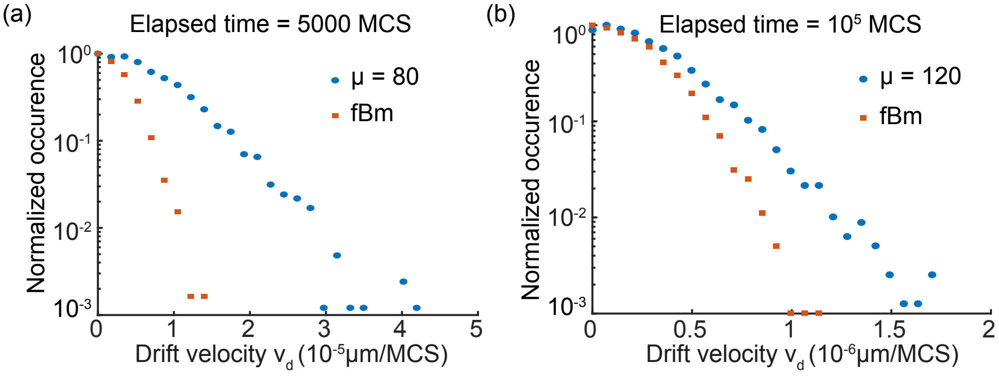


**Fig. S5.** (a) Drift velocity distribution comparisons as in Fig.3(b,c), but at shorter elasped time of 5000 MCS. The differences between bridging simulations and fBm model is amplified. Parameters. $\mu = 80, \lambda = 10^5$ MCS (b) Same as in (a) but for a higher level of bridging. The overall tails get smaller and the difference between fBm and bridging simulations increases as compared to Fig. 3(d). Parameters. $\mu = 120, \lambda = 80000$ MCS.

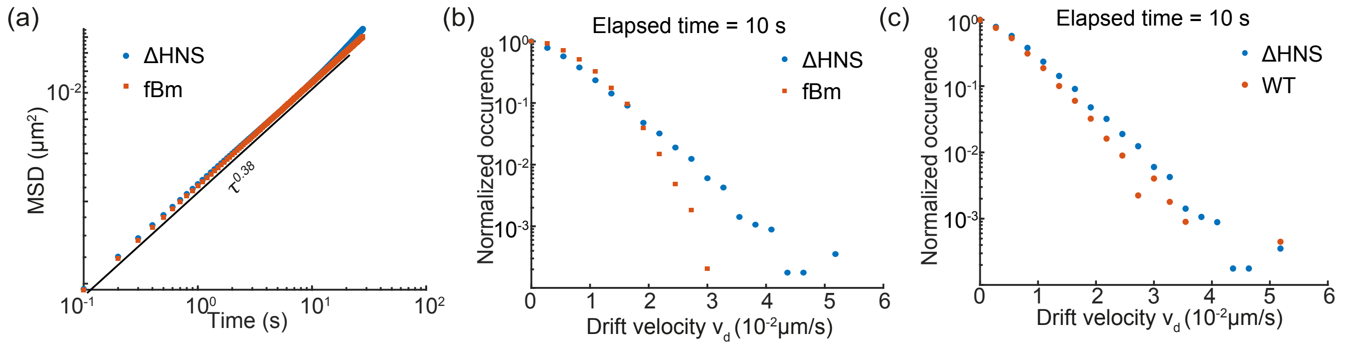


**Fig. S6.** (a) Ensemble averaged MSD from FBM simulations overlaid with ΔH-NS data (Fit, $0.0009\tau^{0.38}$). (b) Drift velocity distributions $v_d$ from FBM simulations have a smaller tail than ΔH-NS data. (c) ΔH-NS has a slightly higher number faster tracks than the wild type.

**RCMs in ΔH-NS.** Similar to the case of the WT and bridging simulations (Fig. 3), parameter matching the mutant ΔH-NS MSD to fBm simulations (Fig. S6(a)) we find that it fails to capture the faster outlier tracks (Fig S10(b)). Note that the mismatch in $v_d$ distribution is not dependent on the power law nature of the mutant MSD curve leading to a better fit. As predicted, we find that the mutant has a slightly broader $v_d$ distribution due to the absence of H-NS (Fig. S6(c)).

#### 3. Bounds on Bridge lifetime

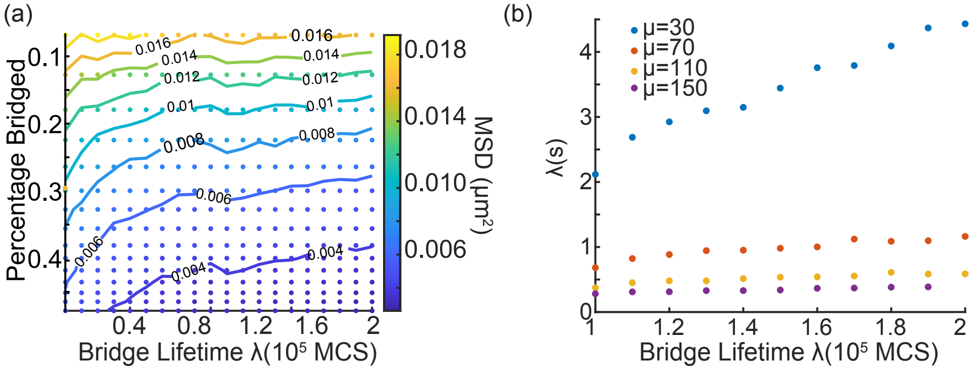


**Fig. S7.** (a) Phase diagram of MSD at 25000 MCS for different parameters. MSD value reached at longer time delays decreases with increasing bridging. (b) Conversion between bridge lifetime $\lambda$ in MCS to seconds. $\lambda$(s) increases linearly with $\lambda$ (MCS) but the slope decreases sharply for higher levels of bridging. Hence, it is computationally challenging to access longer bridge lifetimes in seconds for higher levels of bridging.

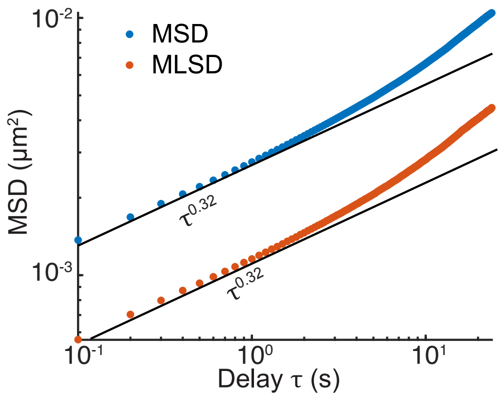


**Fig. S8.** Mean logarithmic squared displacement (MLSD) also shows a transition to higher exponent at longer delays.

**Variability in $\alpha$ and transition in MSD curve.** Time Averaged MSD of a single particle suffers from high variation and random errors. A possible way to mitigate this is by studying the Ensemble Averaged MSD. While this is generally adequate for estimating various diffusion parameters, it has been argued that it suffers from inaccuracies if the underlying population has heterogeneous sub-diffusion. We can calculate the Mean Logarithmic Squared Displacement (MLSD) to account for the exponential dependence on delays (12),

$$r_l^2(\tau) = \log\left(\sum_{t=1}^{T-\tau}[r_n(t+\tau) - r_n(\tau)]^2\right) \quad [3]$$

We an effective exponent $\mu$,

$$\langle r_l^2(\tau)\rangle = \langle \log(D)\rangle + \mu \log(\tau) \quad [4]$$

Analysing our experimental trajectories of WT *E.coli* cells, this procedure did not produce any significant differences in our MSD curves (Fig. S8). While it does not exclude other effects like photo-bleaching, the underlying heterogeneity in $\alpha$ at different time lags might be a real effect arising from a sub-population of trajectories with more mobility (RCMs).

#### 4. Loci tracking experiments

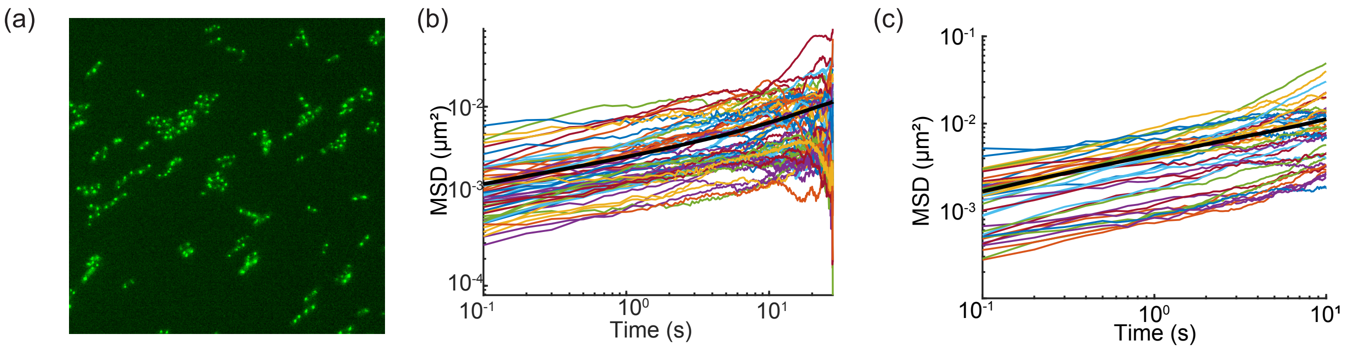


**Fig. S9.** (a) Snapshot of microscopy experiment of GFP-ParB/**parS** labelled *ori* loci in *E.coli.* (b) Sample MSD curves of individual foci obtained from WT strain (set 1). (c) Sample MSD curves of mutant ΔHNS (set 1). Ensemble averaged MSD is represented by the black lines (b,c).

| Scaling exponent $\alpha$ | | | | | | | | | | | | | |
|---|---|---|---|---|---|---|---|---|---|---|---|---|---|
| Strain | set 1 | Tracks | set 2 | Tracks | set 3 | Tracks | set 4 | Tracks | set 5 | Tracks | set 6 | Tracks | All |
| WT | 0.31 | 3952 | 0.319 | 525 | 0.328 | 1193 | 0.339 | 477 | 0.3325 | 506 | — | — | 0.321 $\pm$ 0.011 |
| ΔH-NS | 0.364 | 12121 | 0.357 | 827 | 0.417 | 1647 | 0.407 | 1682 | 0.378 | 1062 | 0.364 | 750 | 0.380 $\pm$ 0.024 |

**Table S1. Table of scaling exponents $\alpha$ observed in experiments from different sessions.**

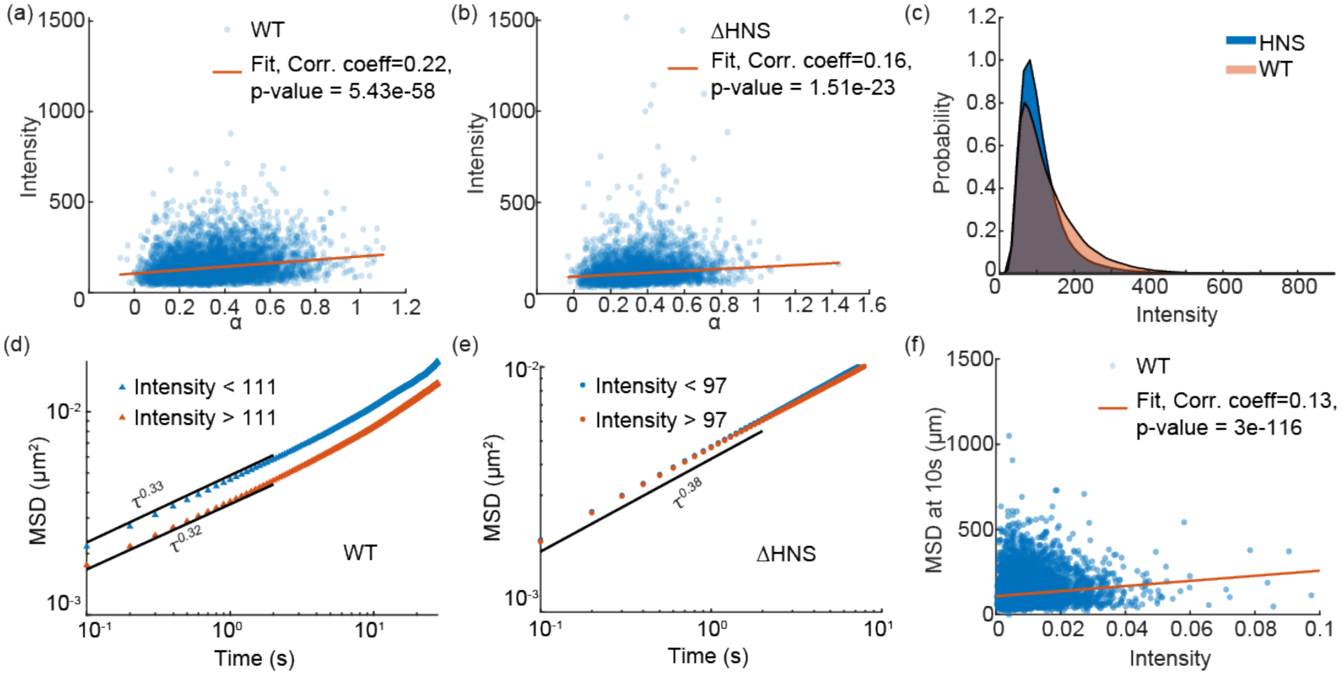


**Fig. S10.** (a) Mean intensity of WT tracks plotted versus scaling exponent $\alpha$ fitted to the entire track length. Shows weak positive correlation. (b) Same as in (a) but for the mutant ΔH-NS. (c) Intensity distributions of WT and ΔH-NS is plotted. WT distribution has a broader tail. (d) Ensemble averaged MSD of WT plotted for two difference sub-populations with intensity $I >$ median$(I)$ and $I <$ median$(I)$. Scaling exponent $\alpha$ shows marginal change while $D_{app}$ decreases with higher intensities (13). (e) Same as in (d) but for the mutant. The sub-populations overlap. (f) Intensity of loci in the WT at a MSD of 10s. We find a very weak positive correlation.

**Intensity of spots does not explain differences in scaling exponents.** We wondered if the difference in MSD scaling exponents between the WT and mutant could be related to the intensity, as it was previously shown that the mobility of loci depends inversely on their intensity (13). In our experimental data, we find that the WT and mutant have comparable intensity distributions and show a very weak correlation between the loci intensity and the scaling exponent $\alpha$ (Fig. S10(a,b)). We also found that the intensity distributions of the strains was comparable, while the WT had a slightly fatter tail (Fig. S10(c)). Comparing the ensemble-averaged MSD of tracks with lower and higher intensity, we found that while intensity affects the apparent diffusion constant $D_{app}$, it has a marginal effect on the scaling exponent $\alpha$ (Fig. S10 (d,e)). The intensity of loci also shows a very weak correlation with the MSD at 10s (Fig. S10(f)). Hence, we conclude that the intensity of loci does not explain the differences between the ensemble-averaged MSD exponents of WT and $\Delta$H-NS.
